# High-resolution Crystal Structures of Transient Intermediates in the Phytochrome Photocycle

**DOI:** 10.1101/2020.09.16.298463

**Authors:** Melissa Carrillo, Suraj Pandey, Juan Sanchez, Moraima Noda, Ishwor Poudyal, Luis Aldama, Tek Narsingh Malla, Elin Claesson, Weixiao Yuan Wahlgren, Denisse Feliz, Vukica Šrajer, Michal Maj, Leticia Castillon, So Iwata, Eriko Nango, Rie Tanaka, Tomoyuki Tanaka, Luo Fangjia, Kensuke Tono, Shigeki Owada, Sebastian Westenhoff, Emina A. Stojković, Marius Schmidt

## Abstract

Phytochromes are red/far-red light photoreceptors in bacteria to plants, which elicit a variety of important physiological responses. They display a reversible photocycle between the resting (dark) Pr state and the light activated Pfr state, in which light signals are received and transduced as structural change through the entire protein to modulate the activity of the protein. It is unknown how the Pr-to-Pfr interconversion occurs as the structure of intermediates remain notoriously elusive. Here, we present short-lived crystal structures of the classical phytochrome from myxobacterium *Stigmatella aurantiaca* captured by an X-ray Free Electron Laser 5 ns and 33ms after light illumination of the Pr state. We observe large structural displacements of the covalently bound bilin chromophore, which trigger a bifurcated signaling pathway. The snapshots show with atomic precision how the signal progresses from the chromophore towards the output domains, explaining how plants, bacteria and fungi sense red light.

## Introduction

Phytochromes are red-light protein photoreceptors, initially discovered in plants^1^ where they regulate essential physiological responses such as shade avoidance and etiolation^2^. With that they are critical to the thriving of all vegetation on earth. Homologous proteins exist in bacteria^3,4^, cyanobacteria^5,6^ and fungi^7^. In photosynthetic bacteria, they regulate the synthesis of light-harvesting complexes^8-12^. In non-photosynthetic bacteria their role is less understood, but they are involved in various processes such as the regulation of carotenoid pigments, which protect from harmful light exposure^4^, in conjugation^13^, plant colonization^14^, quorum sensing and multicellular fruiting body formation^15,16^. Bacteriophytochromes (BphP) have also been successfully used as infrared fluorescent tissue markers in mammals^17^.

Phytochromes consist of 2 modules, where the N-terminal photosensory core module (PCM) is attached to a C-terminal effector module^18^. The latter module provides enzymatic activity together with a so-called N-terminal extension^19^. In plant and class I BphPs the PCM consists of three domains called PAS (Per-ARNT-Sim), GAF (cGMP phosphodiesterase/ adenylate cyclase/FhlA), and PHY (phytochrome-specific) (Fig. 1 a, b). The module is conserved from bacteria to plants and holds a covalently bound bilin chromophore, an open chain tetrapyrrole, which is biliverdin IXα (BV) in bacteria (Fig. 2 b). Hallmark features are a conserved water molecule in the center of the biliverdin, called the pyrrole water (PW)^20^, the so-called PHY (sensory) tongue, which changes fold in the Pr-to-Pfr transition^21^, and the long helix along the dimer interface, which spans the entire PAS/GAF and PHY domains^22,23^. The C-terminal effector domain is divergent between species and is often a histidine kinase in bacteriophytochromes^18,24,25^ Full length phytochromes are difficult to crystallize, but the PCM forms crystals that diffract to 2 Å resolution and beyond^26^. They are particularly suited for time-resolved crystallographic investigations.

**Figure 1.**
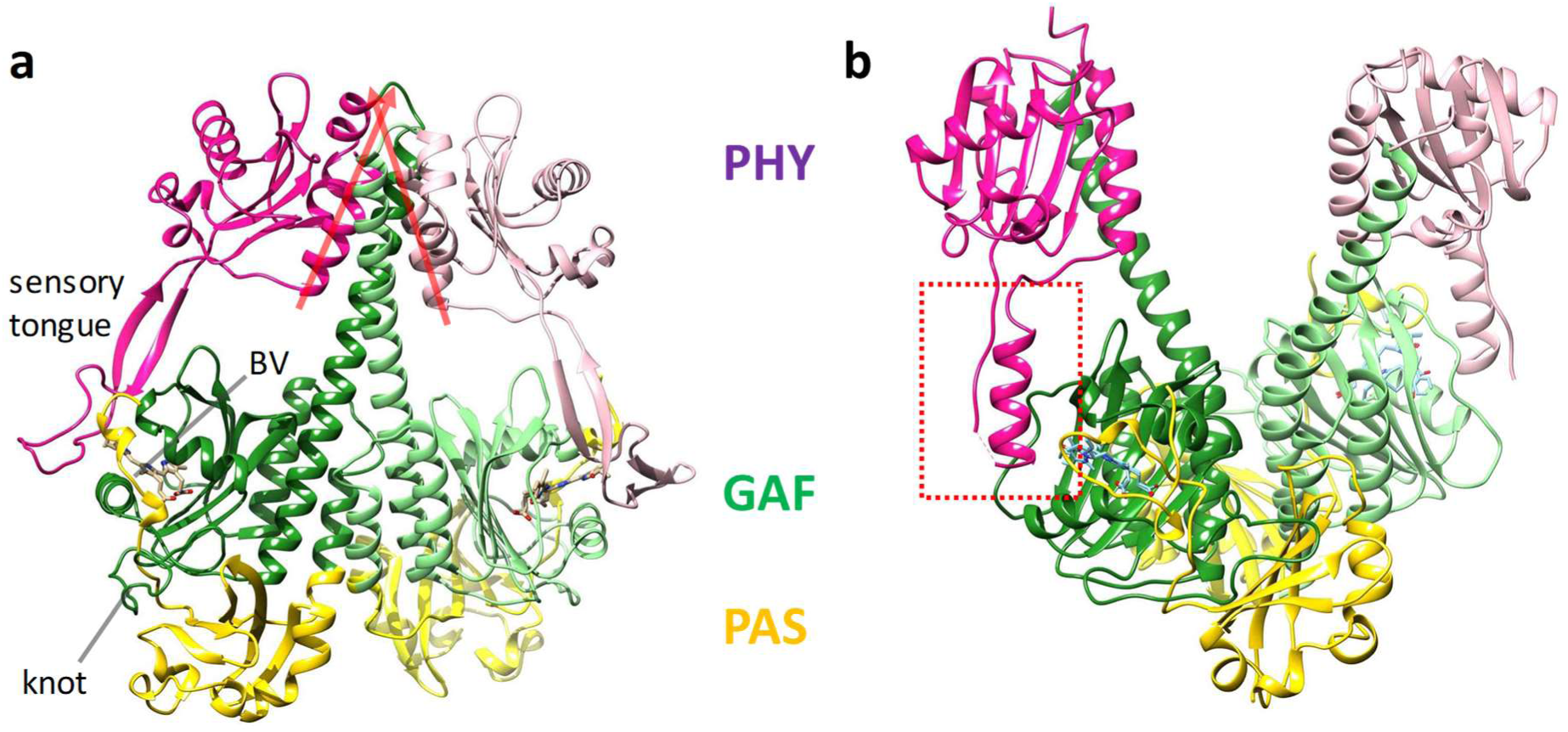
Phytochrome structures. PAS, GAF and PHY domains are displayed in yellow, green and magenta. (a) The *Sa*BphP2 PCM structure in the Pr state, the directions of the short N-terminal helices are shown by red arrows. The biliverdin (BV) chromophore, the characteristic knot, and the sensory tongue are marked. (b) the (static) *Dr*BphP PCM structure in the Pfr state. The sensory tongue of the PHY domain is highlighted with the red-dashed box.

**Figure 2.**
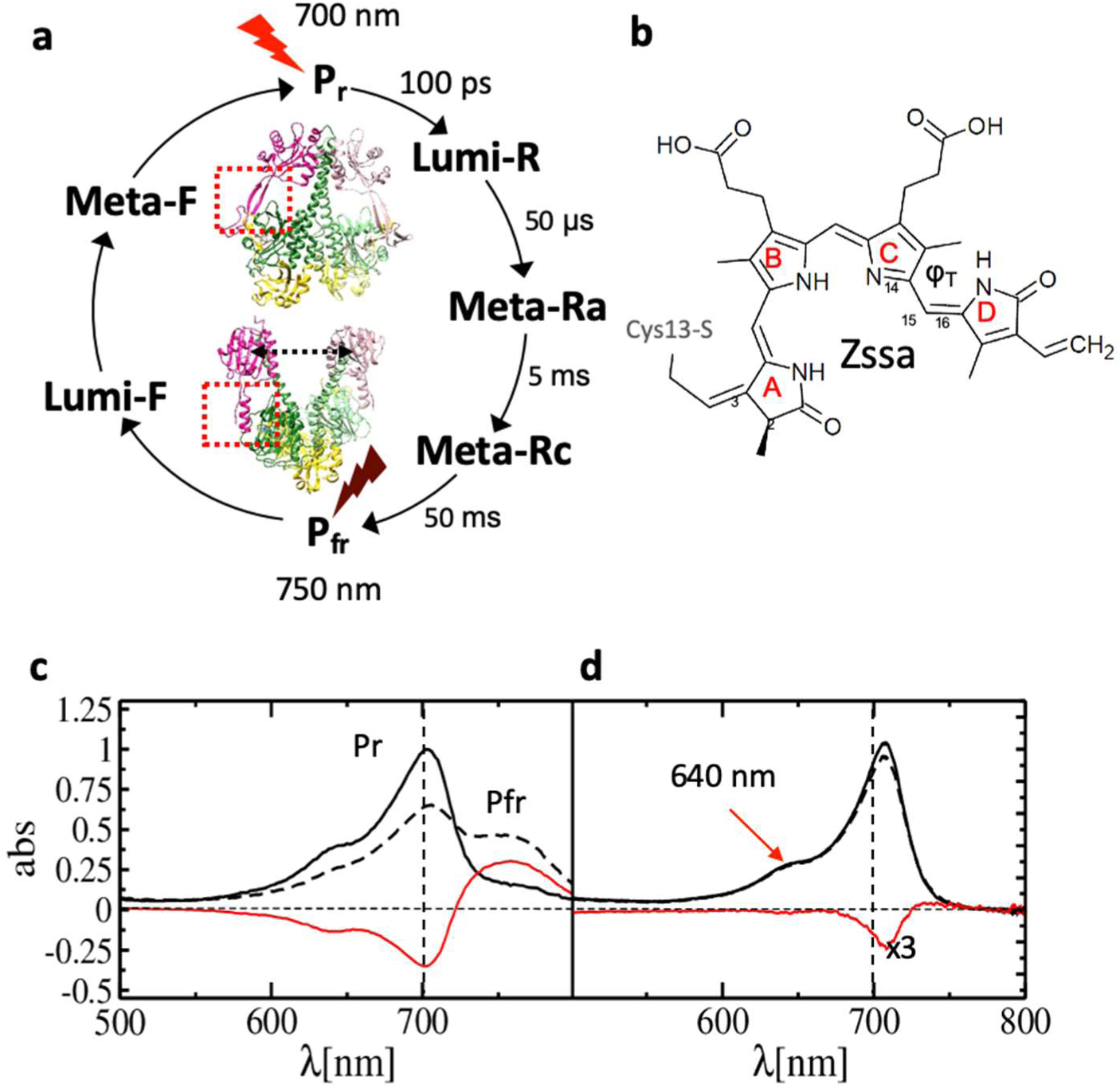
Phytochrome Photocycle, and UV-vis absorption spectra of *Sa*BphP2. (a) Phytochrome photocycle, approximate time scales for intermediate (Lumi-R, Meta-Ra and Meta-Rc) formation are shown. Pfr is formed after about 50 ms. The two half cycles can be driven by illuminating the stable Pr and Pfr states (displayed for the *Sa*BphP2 and *Dr*BphP PCMs, respectively) by red and far red light. (b) The chemical structure of BV bound to Cys-13 in the phytochrome. The torsional angle fT that defines isomerization/rotation about the double bond Δ15,16 is marked. In Pr the structure is all-*Z* syn-syn-anti. (b) *Static* absorption spectra of the *Sa*BphP2 Pr (solid line) to Pfr (dashed line) transition in solution. The difference is shown in red. (c) As (b) but in the crystal. The transition is initiated by 640 nm LED light (arrow). The difference (red line) is enhanced 3-fold.

Phytochromes display a photocycle (Fig. 2 a) with two half-cycles that are driven by two different wavelengths of light. In classical phytochromes, the dark-adapted state, denoted as Pr, absorbs red light (λ ∼700 nm), which causes a *Z* to *E* isomerization of the C15=C16 double bond within its bilin chromophore. Subsequent conformational changes of the entire protein end in a far-red light absorbing state, denoted as Pfr. The Pfr state either relaxes thermally back to Pr, or can be driven back to Pr by far-red light (λ ∼750 nm). The structural changes associated with the Pr to Pfr transition modulate the enzymatic activity of the phytochrome^27,28^. Although the Pr and Pfr states have been structurally characterized in detail using the *Deinococcus radiodurans (Dr)*BphP PCM^21-23,29^, structures of the nanosecond intermediates Lumi-R and Lumi-F as well as those of the longer-lived intermediates in each photo-halfcycle (Fig. 2 a) are missing. In the Lumi-R intermediate, the BV chromophore is in the electronic ground state. The 15*Z* anti (Fig. 2 b) to 15*E* anti isomerization of the C15=C16 double bond between rings C and D of the BV chromophore should have taken place, resulting in a nearly 180° rotation of the D-ring^30-32^. (Fig. 2 a).

Through the latest developments in time-resolved serial x-ray crystallography (TR-SFX), the 1 ps structure of the truncated *Dr*BphP chromophore binding domain (CBD) that consists only of the PAS and GAF domains was determined^33^. 1ps after photoexcitation the BV D-ring in the *Dr*BphP CBD rotates counter-clockwise, while the PW becomes photodissociated from the chromophore binding pocket. Displacements of important, conserved amino acid residues are observed already at 1 ps. For example, the conserved Asp-207 in the GAF domain moves significantly, which could imply signaling directed towards the PHY-sensory tongue. However, the PHY domain is not present in the CBD construct. Experiments on the entire PCM including the critical PHY domain and sensory tongue are necessary to understand how the light signal is transduced to the C-terminal enzymatic domain.

Previous attempts to initiate the photocycle in PCM crystals of various BphPs at room temperature were unsuccessful presumably because the PCM constructs were not photoactive in the crystal form, the illumination protocol was sub-optimal and/or the spatial resolution reached at room temperature was not sufficient^16,34^. Recently, we published the structure of a classical phytochrome from non-photosynthetic myxobacterium *S. aurantiaca*, denoted *Sa*BphP2 PCM solved to a resolution of 1.65 Å at cryogenic temperatures (100 K) in the Pr form. *Sa*BphP2 PCM microcrystals are photoactive (Fig. 2 c) and diffract to 2.1 Å resolution at room temperature^26^ which provides an opportunity to describe the Pr to Pfr transition by TR-SFX experiments.

The TR-SFX experiments on the *Sa*BphP2 PCM reported here were conducted at the Japanese XFEL, the Spring-8 Angstrom Compact X-ray Laser (SACLA). They resulted in room temperature structures 5 ns and 33 ms after light illumination of the Pr (dark) state with 640 nm laser pulses (Methods and Extended Data Tab. 1). Results are discussed in terms of extensive rearrangements of BV, specifically the D-ring, the PW and neighboring water network and conserved amino acids in the GAF and PHY domains.

## Results

### Difference Electron Density at 5ns and 33 ms

The difference electron density (DED) maps calculated at the 5 ns and 33 ms time delays shows a large number of correlated positive and negative DED features in the PAS-GAF as well as in the PHY domains (Fig. 3, Extended Data Tab. 2, see Methods and Extended Data Table 3 for a statistical assessment). These features indicate structural changes through the entire *Sa*BphP2 PCM dimer. The control map at 66 ms only contains spurious features, supporting this assignment (Fig. 3). In all previous ns time-resolved crystallographic experiments on photoactive yellow protein^35,36^, myoglobin^37,38^, and others ^39,40^DED features are mostly localized to the chromophore and a few residues. As a consequence, the DED map sigma level is determined by the noise in the DED map as has been shown previously^41^. Here, this is different. The map sigma level is determined by both the noise and the signal. The large number of difference features poses a formidable challenge for the interpretation of the DED maps, as well as for structure determination. The features must be interpreted locally near the chromophore and the chromophore pocket, and more globally for the entire *Sa*BphP2 PCM dimer (see Methods). Standard deviations (σ) of the DED maps are determined by both the noise and the signal. We therefore use the σ values of the 66 ms control DED map to contour the 5ns and 33ms maps and to identify chemically meaningful signals.

**Figure 3.**
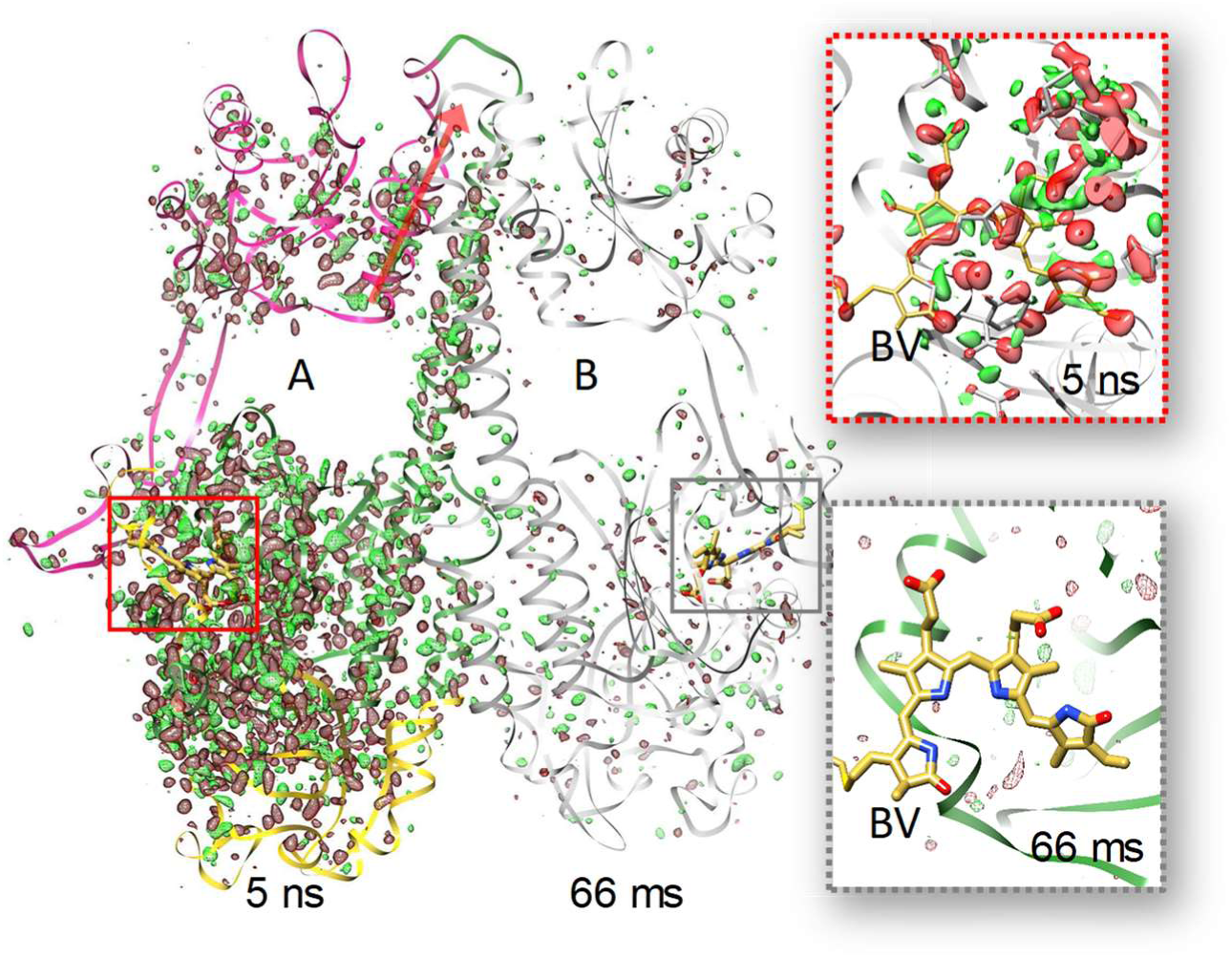
Difference maps for 5 ns and 66 ms time delays overlaid on the *Sa*BphP2 PCM dimer. The 5 ns DED map is displayed with subunit A, the 66 ms map on subunit B (gray). Contour levels: red −3σ, green 3σ. PAS, GAF and PHY domains of subunit A in yellow, green, magenta. Red arrow: direction of the C-terminal helix that connect to the coiled-coil linker in the full length BphP. Note the numerous DED features along the dimer interface helix into the PHY domain and the C-terminal helices at 5 ns. Inserts show the corresponding difference maps in the biliverdin (BV) binding pocket. At 66 ms, only spurious, random DED features are present. The chromophore is essentially free of signal.

### Ring-D Orientations at 5ns and 33ms

Substantial DED features are observed on and near the BV chromophore (Fig. 3, 5 ns insert and Fig. 4). Strong negative DED features on the D-ring carbonyl, methyl and vinyl mark substantial structural rearrangements in both subunits. Interestingly, positive features that identify D-ring orientations upon light illumination differ in subunit A and in B. In subunit A, there are strong lateral features (β_1_ to β_4_) that support a clockwise ∼90° twist of the D-ring (Extended Data Fig. 1 b-e) when viewed along the chromophore axis from the D to the A-ring. These features can be reproduced by calculated difference maps (compare Extended Data Fig. 1 b,c and d,e). In addition, positive features ξ (Fig. 4 a) are oriented in a way that support a 180° rotation. For the interpretation of features ξ, extrapolated electron density (EED) maps were necessary since the strong negative density on the D-ring carbonyl tends to eliminate close-by positive features. Ring-like electron density appears in the EED maps (Fig. 4 b) indicating a fully isomerized D-ring. Accordingly, two conformations of the chromophore are need to interpret the positive DED to completion, a ∼90° clockwise D-ring twist and a fully isomerized ∼180° clockwise D-ring rotation (Fig. 4 b, Extended Data Tab. 4).

**Figure 4.**
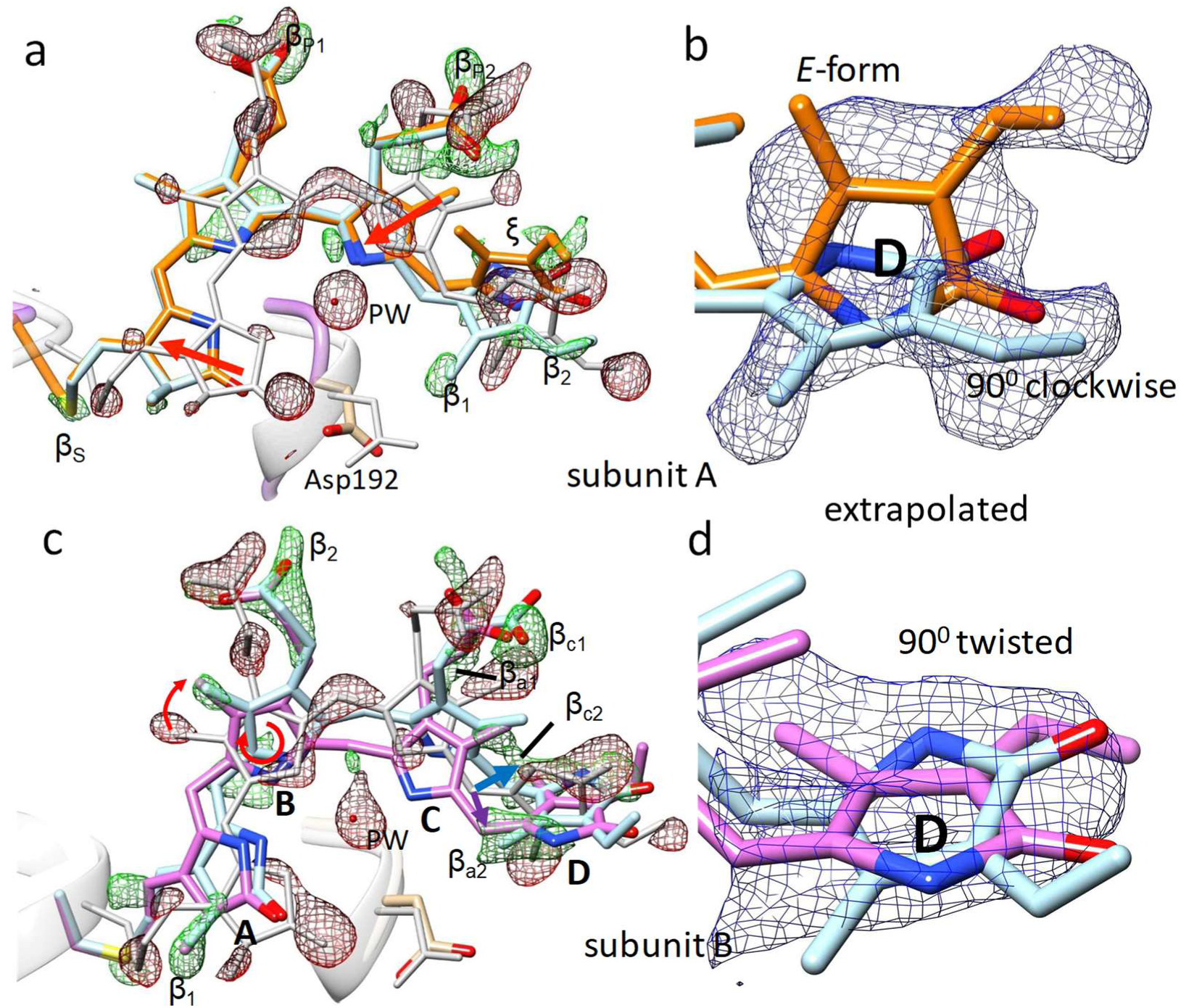
Chromophore displacements and D-ring rotations. DED is displayed in red and green, contour levels: red, −2.7 σ (- 3 σ of DED_66ms_), green, 2.7 σ (3 σ of DED_66ms_), EED is shown in blue with N_c_ = 22 (b) and N_c_ = 19 (d), contour level 1.5 σ. (a) overall chromophore configuration in subunit A at the 33 ms time delay. Gray: reference (dark), orange: intermediate at 33 ms. subunit A, reference structure in gray, 90° clockwise D-ring rotation (light blue), full isomerization (orange). Chromophore slides in the direction of the red arrows. (b) EED (blue), N_c_ = 22, on the D-ring in subunit A, colors as in (a). (c) Chromophore configuration in subunit B at 5 ns. Positive features β determine the ring positions. Feature β_a_ (behind the chromophore plane) enforce a C-ring tilt resulting in a counter-clockwise rotation (purple and purple arrow). Clockwise rotation (blue arrow) shown in light blue. (d) EED (blue), N_c_ = 19, on the D-ring in subunit B, colors as in (c).

**Figure 5.**
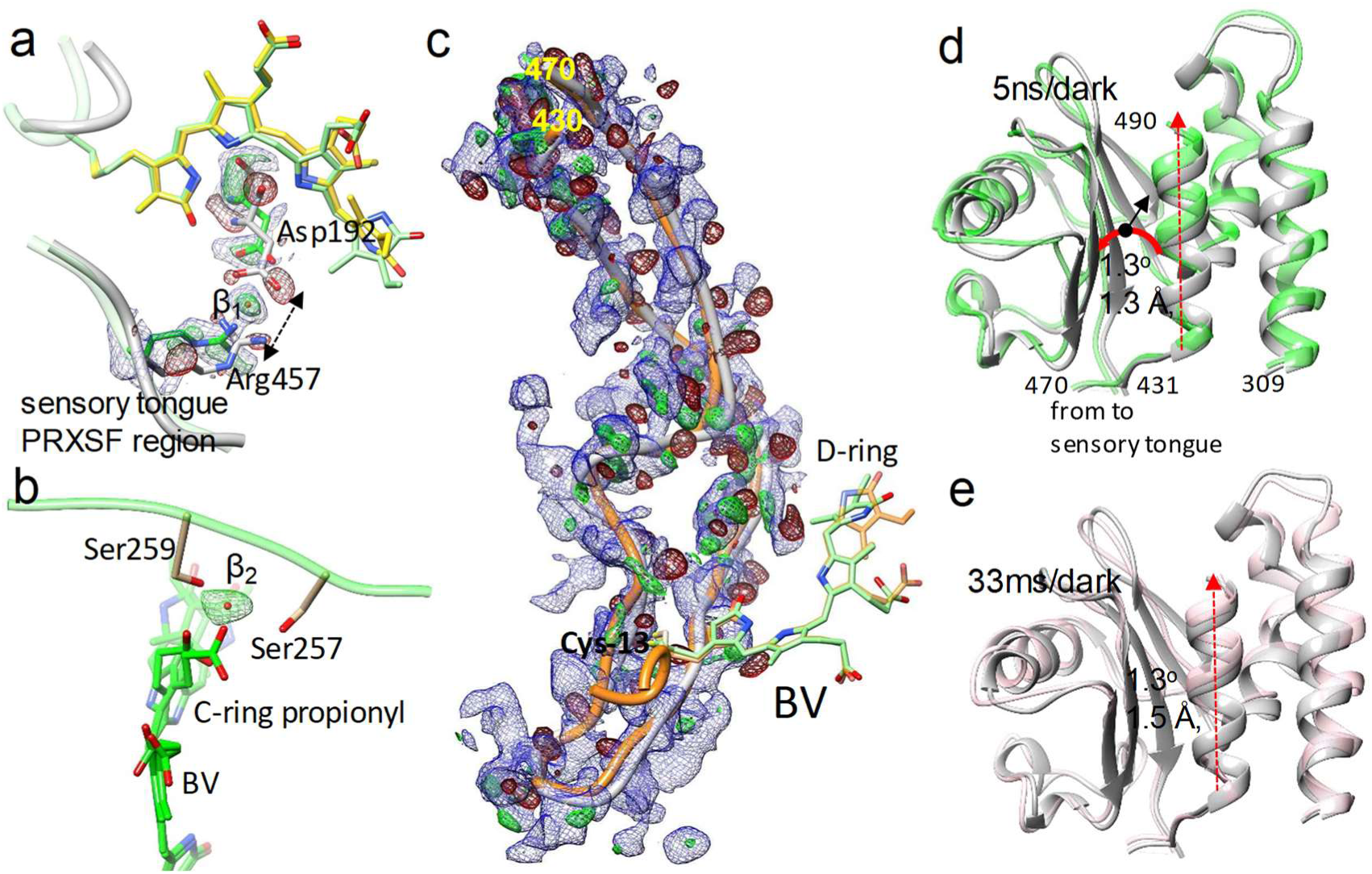
Local and global structural changes. DED in red and green (-/+ 3σ contour), EED in blue (1.2 σ contour). (a) Separation of the sensory tongue from the BV binding region (dotted arrow). Asp-192 and Arg-457 are marked. The BV chromophore with the 90° twisted and fully isomerized D-ring is shown in yellow and green, respectively. β_1_, positive DED feature interpreted by a water molecule. (b) The BV (green) C-ring propionyl detaches from Ser257 and Ser259 which coordinate a water (feature β_2_, green positive DED) instead. The structure is displayed for subunits B (33 ms) where the D-ring twists ∼90° both clockwise and counter-clockwise. (c) The sensory tongue region. Gray: structure of the reference state, orange: structure at 33 ms. Residues at the beginning and the end of the region are marked. The chromophore is shown with the twisted D-ring (green) and the fully isomerized form (orange). (d) The PHY domain region. Comparison of the 5ns structure (green) to the reference structure (gray). Sequence numbers are marked. The PHY domain centroid (black dot) moves by 1.3 Å (black arrow), and rotates (red curved arrow) by 1.3°. The connection to and from the sensory tongue is marked. (e) PHY domain at 33 ms versus reference. Similar displacements as in (a) are observed. Displacement of the c-terminal helix is marked by the red dashed arrow in (d) and (e).

In subunit B, features ξ are absent (Fig. 4 c). In accordance, EED maps (Fig. 4 d) do not support a fully isomerized configuration (as in subunit A) for both the 5 ns and the 33 ms time delays. Instead, strong positive features determine the geometry of the BV A to C-rings; see Fig. 4 a and c for a comparison of the DED in both subunits. In subunit B the entire BV pivots about the B-ring (Fig. 4 c) which leads to strong C-ring and D-ring displacements. To interpret positive features β_c1_ and β_c2_ (‘c’ for clockwise) C-ring must be tilted backwards (blue arrow in Fig. 4 c) and the ring-D can only be oriented clockwise (light blue BV structure in Fig. 4 c,d) to fit the DED. This leaves a strong feature β_a1_ which is located behind the C-ring plane. To reproduce this feature, the ring C propionyl must tilt in the opposite direction (purple arrow in Fig. 4 c) which leads to a displacement of carbon atom C_15_ forward. Then, the counter-clockwise D-ring orientation (in pink) fits the DED.

### Amino acid and water network rearrangement in the chromophore binding pocket and the sensory tongue

Strong negative DED features indicate that the PW photo-dissociated from BV in both subunits at 5ns and 33ms (Fig. 4, Extended Data Tab. 2). Moreover, significant displacements of the conserved Asp-192 of the PASDIP consensus sequence in the GAF domain and the Arg-457 of the PRXSF motif^22,42^ in the PHY domain are observed in subunit A (and at 33 ms also in subunit B). Asp-192 and Arg-457 form a salt bridge, anchoring the PHY tongue to the chromophore region in the Pr state. This connection is broken at 5ns. A strong positive DED feature between these two amino acids is observed, indicating a water molecule (Fig. 5 a). Furthermore, the conserved Tyr-248 in proximity to Asp-192 adopts a dual conformation at 5 ns and 33 ms (Extended Data Fig. 2). Similarly, a dual conformation is observed for the conserved His-275 (at 33 ms) that forms a hydrogen bond to the D-ring carbonyl in the Pr state of the *Sa*BphP2 and other classical BphPs. In contrast to these amino acid rearrangements, the structure of the sensory tongue is only locally affected (Fig. 5 c). A β-sheet to α-helix transition is not observed which coincides with only minor changes of the PHY domain position (Fig. 5 d,e).

On the other side of the chromophore, opposite to the tongue region, strong features in the DED maps at 5ns and 33 ms indicate that the BV C-ring propionyl drags the conserved Ser-257 and Ser-259 along at 5ns and 33 ms. (Fig. 5 b, see also Extended Data Tabs. 2 and 4). A positive DED feature in the B subunit indicates the appearance of a water molecule that may form hydrogen bonds with the C-ring propionyl as well as with Ser-257 and Ser-259. Moreover, the DED maps show correlated negative and positive features within the PAS-GAF domains and along the long helices that form the dimer interface (Fig 3), pushing outwards the C-terminal helix that connects to the output module (Fig 5 d and e). We propose that the structural changes in the chromophore pocket initiate the signal that is transduced along the long helices ‘wiring’ together the BV chromophore and effector domains.

## Discussion

### Global and local structural relaxations at 5 ns and 33 ms post illumination

In the *Dr*BphP CBD protein relaxations can already observed at 1 ps and 10 ps^33^. Although individual BV ring displacements are observed, the chromophore essentially stays at the position that it also occupies in the dark structure (Extended Data Tab. 5). Despite this, the signal has penetrated deep into the BV-pocket of the GAF domain^33^. On the ns time-scale the signal is expanding further through the entire PCM of the classical *Sa*BphP2. In contrast to smaller proteins, such as PYP and myoglobin, the *Sa*BphP2 is large and flexible and can react readily and fast to local chromophore perturbations. In the *Sa*BphP2 PCM structures shown here, protein relaxations are advanced enough that large chromophore displacements are observed in both subunits. Especially in subunit B, large chromophore geometry distortions are present (Extended Data Tab. 4) as the C-ring tilts out in both directions. It appears as if the energy of the absorbed photon is stored in a distorted BV geometry that drives protein relaxations. Chromophore geometry distortions are also found in early intermediates of unrelated proteins such as the PYP^35^. Distortions of the BV chromophore of the phytochrome photocycle have been also predicted by time-resolved spectroscopy^31^. They were directly observed in temperature scan cryo-crystallography experiments performed on the bathy phytochrome *Pa*BphP from *Pseudomonas aeruginosa*^43^ (Extended Data Tab. 5) further suggesting that distorted chromophore conformations are part of an important mechanism to advance photochemical reactions. As the BV chromophore position changes, it strongly affects amino acid residues near the BV propionyl moieties that shift in unison with the chromophore. Examples are listed in Extended Data Tab. 2 and 4, and are discussed further down. The species with the full 180° rotation of the D-ring might be associated with a key Lumi-R like intermediate in the phytochrome photocycle (Fig. 2 a). To determine the specific time point where the 180° rotation begins, additional data collected at different time delays are required.

The absence of the fully isomerized D-ring isoform in subunit B can likely be explained by differences in the subunits related by non-crystallographic symmetry. As the structure of the sensory tongue is essentially identical in both subunits, it is unlikely the reason for this behavior. By inspecting the region near Cys-13 to which the BV chromophore is bound, differences are found. Distances to symmetry related molecules are different for subunit A and B. In subunit A the distances from Arg-15 to Gln-139 and Cys-13 (S) to Lys-136 (N_z_) (Gln-139 and Lys-136 belong to the molecule related by crystallographic symmetry) are 6.0 Å and 7.2 Å, respectively. These distances are smaller in subunit B (5.2 Å, and 4.0 Å, respectively). These differences likely have an impact on chromophore relaxations, as in subunit B BV structure appears more distorted than in subunit A (see twisting angles for subunits A and B in Extended Data Tab. 4).

### Rearrangement of water network and neighboring amino acids

The PW forms a stable hydrogen bond network with BV rings A-C in both the Pr and Pfr states, but it is absent at 5 ns and 33 ms. It photo-dissociates already within 1 ps in the *Dr*BphP CBD fragement^33^. While the twisting motion of D-ring has been the working model for phytochrome activation and now has been confirmed, the disappearance of the PW is surprising. Given the large sliding motions of the chromophore (Fig. 4 a and c), this now makes sense as the absence of the PW most likely enables these displacements. It is interesting to note that the PW dissociates very early^33^ and rebinds back to BV in Pfr^21,29^. The PW may have a dual role in facilitating the structural transition and stabilizing the reaction product (Pr as well as Pfr) in both halves of the reaction cycle. Both, the rotation of the D-ring together with photodissociation of the PW likely are the main triggers for subsequent protein structural changes. Together, they transduce the light signal to the sensory tongue of the PHY domain and cause relaxations of the GAF domain that propagate further up the long helices along the dimer interface.

### Sensory tongue and the PHY domain

The sensory tongue connects the PHY domain directly with the chromophore region (Fig. 5). During the full Pr the Pfr transition the sensory tongue undergoes extensive structural transitions from a beta sheet to an alpha helix^21,29^. In the Pfr state, Pro-456 becomes adjacent to the D-ring and forms a hydrogen bond with Tyr-248. This is stabilizing the D-ring in E configuration. It is interesting to ask how this shift is initiated. In the present microcrystals, the transition is not observed (Fig. 5 c). We ascribe this to the crowded environment of the crystals. However, the tight Asp-192 to Arg-457 salt bridge is already broken at 5 ns and a water is inserted in between the residues. This is an important first step to enable the sensory tongue to rearrange. We therefore conclude that the signal is transduced to the PHY tongue via the displacement of the chromophore that enforces the movement of Asp-129 and the photo-dissociation of the PW.

### Propagation of the light signal

Caused by D-ring rotation, the conserved Tyr-248 moves (Extended Data Fig. 2). This destabilizes interactions with the neighboring amino acids and the water network. Arg-457 and Aps-192 form new hydrogen bonds with a water molecule (Fig. 5 a). As the chromophore slides substantially (Fig. 4), it induces structural changes in the GAF domain sensed by the conserved serines 257, 259 and 261 and multiple other amino acids near the chromophore. His-275 loses contact with the ring-D carbonyl (distance: > 4 Å) and with the more distant Arg-157 (now ∼4.0 Å) that lead to substantial GAF domain relaxations which are ultimately relayed to the PHY domain through the long dimer-interface helices. As the speed of sound in protein crystals is about 2000 m/s^44^, heat expansion through 100 Å of protein (roughly the length of the *Sa*BphP2 PCM) in 5 ns cannot be excluded. However, relaxations at 33 ms are very similar to those at 5 ns. Heat produced locally after chromophore light absorption should have dissipated by then, and the DED features at 5 ns rather represent genuine protein relaxations.

The changes on the long dimer helix and the C-terminal helix (Fig. 3) suggest a mechanism of signal transduction that does not rely exclusively on the opening of the PHY domains (Fig. 1 b). It seems as the long helices, and not so much the sensory tongues, translate the signal towards the small C-terminal helices that are connected to the coiled-coil linker region of the effector domain (Fig. 5 d,e, red arrows). This confirms a suggestion that was made based on the static crystal structures of the bathy *Pa*BphP PCM^23^ and compares favorably to signal transduction in transmembrane sensory proteins^45^. Only small PHY domain displacements (Fig. 5 d,e) are necessary for signal transduction. Since the linker helices of the sister monomers are at an angle, translations along their axes will slightly change the relative orientation of the effector domains, and hence their activity, possibly modulated by a shift in register of the coiled coil linker ^27^.

### Summary and outlook

The short-lived structural intermediates presented here establish that it is indeed the isomerization of the D-ring, which drives the photoconversion in phytochromes. The remaining rings of the chromophore move notable distances and the movements are heterogeneous between the different subunits. The presence of unproductive BV conformations, may explain the relatively low quantum yield for the Pr to Pfr transition (approximately 10-15%). Nevertheless, we observe a fully isomerized BV configuration and establish that photodissociation of the PW and the displacements of the strictly conserved Asp-195 and Tyr 248 lead to a disconnection of the PHY sensory tongue from the chromophore region. Finally, the data show strong evidence for a structural change along the long helices at the dimer interfaces that transduce the signal further towards the PHY domains.

Earlier time points within the *Sa*BphP2-WT PCM photocycle should be collected to assess when the large BV chromophore displacements begin. Large scale structural changes are limited by the crystal packing. Therefore, methods, which act on proteins in solution should be explored to make further progress. Recently, solution NMR spectra of a full PCM was assigned for a phytochrome^46^ and new developments in cryoEM^47^ bring atomic resolution of macromolecular structures within reach without the need for crystals. Calculations are underway^48^ explaining how to obtain structures from single biological macromolecules, such as the full-length, intact BphPs at XFELs.

## Methods

### Protein purification and crystallization

Microcrystals of the *Sa*BphP2-PCM were grown as described^49^ by mixing a mother liquor consisting of 0.17 M Ammonium acetate, 0.085 M Sodium citrate tribasic dihydrate pH 5.6, 25.5% w/v Polyethylene glycol 4000, 15% v/v Glycerol (cryo-screen solution) and 3 % w/v Benzamidine Hydrochloride, with 60 mg/mL protein (3:2 protein to mother liquor ratio). The mixture was seeded with finely crushed macrocrystals. After 4 days, the microcrystals were collected and concentrated to about 10^11^ crystals /ml and subsequently folded into a tenfold amount of nuclear grade grease^50,51^. All steps in crystallization and tray observations were performed under green safety light.

### Experimental Setup

Pump-probe experiments were conducted at beamline BL2 at SACLA using a nanosecond laser^52,53^. For our nanosecond TR-SFX experiments a two-sided laser illumination geometry was used where a split laser beam intercepts the X-rays and the viscous jet at a 90° angle from opposite sides. A relatively large laser fluence of 3.5 mJ/mm^2^ was chosen for each side, respectively. The laser fluence was chosen based on absorption measurements on grease crystal mixtures (Extended Data Fig. 3, see also below). For femtosecond TR-SFX experiments X-rays and laser illumination are parallel ^33,39,54-57^. Then, the effective ‘hit-rate’ of the laser illumination is equivalent to the X-ray hit-rate ^54,55^ and shading by other crystals in the viscous jet does not play a role. This has consequences for the selection of the laser fluence, especially for fs laser illumination, which are discussed ^54,55^. In contrast, for a perpendicular geometry as employed here, the entire path of the X-ray beam through the crystal must be illuminated by the laser. This leads to an effective laser beam size that is much larger than the X-ray beam (Extended Data Fig. 3 b). The large effective laser beam size is likely intercepted by other crystals in the relatively thick (100 μm) viscous jet. This results in substantial shading by crystals not exposed to the X-rays. To roughly estimate this shading, the crystal-grease mixtures were sandwiched between cover slides kept apart by 50 μm washers to match the optical path through half of the 100 μm thick viscous jet. Absorption was measured with a microscopectrophotometer located at BioCARS (APS, Argonne National Laboratory). Grease mixed with *Sa*BphP2 PCM microcrystals shows an absorption of 0.75 at 640 nm (Extended Data Fig. 3 a), which corresponds to a 5 fold reduction of the incident fluence. In addition, at 640 nm the absorption is only 40% of that at the maximum at 700 nm. Accordingly, a 3.5 mJ/mm^2^ fluence at 640 nm is equivalent to only about 0.28 mJ/mm^2^ at the absorption maximum. Because of the 2-sided illumination the total fluence at a crystal probed in the middle of the jet is 0.56 mJ/mm^2^ (with reference to the absorption maximum). Given the previous experiences with photoactive yellow protein^36,56^, 0.56 mJ/mm^2^ is well below the threshold to generate damage even with femtosecond laser pulses ^58^, and does not play any adverse role with nanosecond laser pulses. We believe that for our experimental geometry strong laser excitation has been essential to boost excitation levels to the extent that analyzable signal is obtained.

### TR-SFX data acquisition and processing

200 µl of the crystal-grease mixture were transferred into an injector reservoir^59^ and extruded into air at ambient temperatures (293 K) through a 100 µm wide nozzle with a flow rate of about 4 μl/min. The photoreaction was started with 5 ns lasers pulses of 640 nm wavelength with a full width half maximum (FWHM) of 52 μm. The laser repetition rate was varied between 15 Hz and 10 Hz (Extended Data Fig. 4). A flow rate of 4 μl/min displaces at least a 300 μm column of grease between the high frequency (15 Hz) laser pulses. 5 ns after the laser pulse the stream of microcrystals was exposed in air to intense X-ray pulses (λ = 1.38 Å) of < 10 fs duration with either 10 Hz or 15 Hz repetition rates. The scattering background was minimized by using a helium-purged collimator. We used a pump-probe, 33 ms, 66 ms (Extended Data Fig. 4 b) data collection strategy to assess whether a once laser illuminated/excited viscous jet volume has left the X-ray interaction region and moved sufficiently that multiple laser excitations of the X-ray probed volume are avoided.

Reference data which are free of laser excitation have been collected previously without the laser^26^. For all experiments, diffraction patterns were collected on a CCD detector with eight modules^60^ and analyzed with a user-friendly data-processing pipeline^61^ consisting of hit-finding with Cheetah^62^, and indexing and Monte Carlo integration by CrystFEL^63^. The hit rate was about 30%. About 50 % of diffraction patterns were successfully indexed. Mosflm and DirAx both were used for indexing. The extracted partial intensities were merged to full reflection intensities using the ‘partialator’ program in CrystFEL. For data statistics, see Extended Data Tab. 1. The full intensities were converted to structure-factor amplitudes by software based on the CCP4 suite of programs ^64^.

### Computation of Difference Electron Density Maps

Weighted difference electron density (DED) maps were calculated as described ^36,65^. The DED maps at 33 ms and 66 ms were inspected for strong DED features near the chromophore. At 33 ms clear signal is present (Extended Data Fig. 5 b). At 66 ms, only spurious and randomly distributed features could be detected (Extended Data Fig. 5 c). This demonstrates that the viscous jet is extruded fast enough to cope with a data collection strategy shown in Extended Data Fig. 4 a (pump-probe, dark) as with this strategy the next laser pulses arrives 66 ms after the previous one. This way, the laser excites a pristine jet volume that is free from contaminations from earlier laser pulses. Data for two time-delays are collected from the same experimental setup, as the first X-ray pulse after laser excitation contributes diffraction patterns for a 5 ns dataset, the second X-ray pulse contributes to a 33 ms dataset. Needless to say, when only one intermittent X-ray pulse is used (Extended Data Fig. 4 a, the reference data must be collected separately with the laser switched off (see above).

### Statistical Analysis of DED Features

The signal content and signal variance (sigma values) in the DED maps were analyzed by histograms. Gaussian fits to the histogram should reproduce the root mean square deviation (RMSD) values of the fft program that calculates the DED when the signal is purely random. When the signal is weak and localized, it only changes the distribution in the flanks of the Gaussian^66^, but not the sigma value. If the signal is strong and everywhere, the fitted Gaussian becomes broader and does not fit the flanks. The noise originates from the experimental error in the difference amplitudes and errors introduced by the Fourier approximation^67^. In the presence of localized signal, the noise and not the signal determines the sigma value of a DED map^66^. Here, this is not the case, and occurs for the first time in time-resolved crystallography. In Extended Data Fig. 6 b, a histogram of DED values derived from difference amplitudes ΔF = F_66ms_-F_Dark_ is shown. The histogram is fit by a Gaussian with a sigma of 0.0125 e^-^/Å^3^. The same value also reported by the fast Fourier program (‘fft’) from the ccp4 suite of programs. The Gaussian fits the histogram perfectly which outlines the random nature of the DED features. A histogram prepared from the 5ns-F_dark_ DED_5ns_ map is shown in Extended Data Fig. 6 a. The Gaussian is broader as in Extended Data Fig. 6 b, and the flanks of the histogram are not fit properly by the Gaussian. If the DED features containing signal would be sparse, the sigma from the fit would be essentially identical to the value obtained from the 66 ms control data only. However, it is larger, 0.0144 e^-^/Å^3^. The form of the histogram is an indication of strong signal throughout the map. For the DED_5ns_ map the sigma value is determined by all of the noise sources described above plus the signal. Consequently, for meaningful comparisons, the DED_5ns_ map must be contoured as a multiple of the sigma value found in the DED_66ms_ control map, as this reflects the error level in a DED map without signal.

As large numbers of DED features were found, it is useful to estimate how many of these features might be generated by the noise sources mentioned above. For this, DED maps were sampled on a 3D grid no larger than 2 (*h*max +1) with h_max_ the maximum h,k and l values within the resolution limit. As an example: at a resolution of 2 Å and a unit cell axis a = 80 Å, h_max_ is 40. Assuming similar values for the other cell axes, the unit cell and its DED content is sampled on a 82 x 82 x 82 grid. This way the DED peaks found in such a map correspond to independent features. Extended Data Tab. 3 contrasts the number of DED features observed at 33 ms to the ones expected to occur randomly in the DED map. On the 3 sigma level, for example, 1839 features are expected to be purely random, and 3167 features are observed. On the 5 sigma level, the probability of a feature to be random is so low that not even one feature is expected in the entire DED map, yet 264 features (132 for a *Sa*BphP2 PCM dimer located in the asymmetric unit) are observed. To bring this into perspective, in the strong difference map determined for the photoactive yellow protein at a 3 ps pump-probe time delay^56^, only 12 features, two per symmetry related PYP molecule, are observed on the 5 sigma level (at 2.1 Å resolution to be compatible with the resolution achieved here, and h_max_ determined as described).

### Structure determination

Structural models were derived from extrapolated maps calculated by adding N*DF to the structure factors calculated from an accurately refined dark state *Sa*BphP2 PCM model. The factor N_C_ required to extrapolate the fraction of excited molecules to 100 % was determined by integrating negative density in the extrapolated maps until the values diverge (Extended Data Fig. 7). For the 5 ns time delay N_C_ is 19 which corresponds to a population of 10.5 % activated molecules in the crystal ^65^. For the 33 ms time delay N_C_ is 22 (about 9% of molecules are activated). The chromophore was moved by hand into the extrapolated maps calculated at 5 ns and 33 ms time delays. The D-ring was rotated about the double bond Δ15,16 to achieve maximum agreement with the DED maps as well as with the extrapolated density. Multiple D-ring orientations were accommodated by generating chromophore double conformations. The apo-SaBphP2 PCM structures were determined by using the real space (stepped) refinement option in ‘coot’. This was followed by a scripted ‘zoned’ refinement, also performed in real space in ‘coot’. For this, the script activates α-helical and β-strain restraints when needed. After the real space refinement the agreement with the difference map was inspected, and if necessary corrected further by hand. Double conformations for certain residues (His275, Tyr 248) were introduced to accommodate DED features that result from the various D-ring orientations. If in doubt, extrapolated maps with higher N (N ∼ 40) were computed to verify the presence or to clarify the absence of individual structural moieties at specific locations in space. A final reciprocal space refinement was conducted using phased extrapolated structure factors (PESF) calculated as described previously^33,65,68^. As calculated structure factors of the reference structure are used to determine the PESFs, the final refinement is biased towards the reference (dark) structure. Therefore, refined differences are (i) real, and (ii) sometimes tend to be underdetermined by a fraction of an Å. Extrapolated amplitudes also amplify errors in the difference amplitudes N_c_ times. Due to this, structures were refined to 2.4 Å which is lower than the resolution limit of the data. Due to the same reasons, occupancies for the chromophore and other amino acid residues double conformations were not refined. Occupancies were rather distributed on equal par among the double conformations.

### Pearson Correlation Coefficient to estimate ring-D orientation

Once a model that interprets the DED is determined, calculated DED maps can be computed from this model and the reference model ^33,68^ by subtracting structure factors calculated from both models. In order to corroborate the assessment of clockwise and counter-clockwise D-ring rotation, the Pearson Correlation Coefficient was used. For the correlation coefficient the observed and calculated DED maps are compared. Since negative DED is always on top of the reference model, the negative DED does not add information to distinguish competing models. Accordingly, only positive DED features near the ring-D region are compared. A Fortran program was developed that reads difference maps in ccp4 format, masks out a specific volume around a pdb-file provided to the program, and calculates the Pearson correlation coefficient (PCC) within this masked volume. Here, the mask volume is determined by the coordinates of the D-ring in both clockwise and counter-clockwise orientations. Within this volume, the PCC was determined by a grid-wise comparison of positive observed and corresponding calculated DED features as:

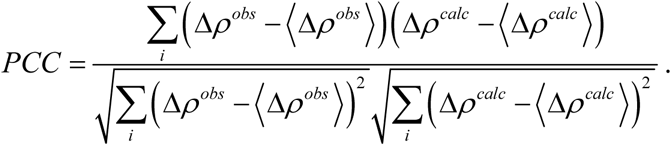

The terms in the bracket are the averages computed from all grid points i in the mask. Although the average DED in an entire difference map is zero, the averages computed here are not zero, since only positive features are evaluated. In addition, map grid points i were selected only when the observed DED values are larger than a certain sigma value. With this, the PCC can be plotted as a function of increasing sigma values (see Extended Data Fig. 8). In subunit A, at 5 ns, only the clockwise rotation is supported in particular by high DED features. In subunit B, the PCC is similar regardless whether a model with a clockwise, a counter-clockwise or a double conformation is examined. This demonstrates that both clockwise and counter-clockwise ring-D rotations produce calculated DED features that explain the observed density equally well. The PCC cannot distinguish between the two ring-D orientations.

### Detailed Views of Structural Moieties

Structural views were generated by UCSF Chimera^69^.

## Acknowledgements

This work was supported by National Science Foundation (NSF) Science and Technology Centers (STC) grant NSF-1231306 (“Biology with X-ray Lasers”). Some results shown are derived from work performed at Argonne National Laboratory, Sector 14 - BioCARS at the Advanced Photon Source. Argonne is operated by UChicago Argonne, LLC, for the U.S. Department of Energy, Office of Biological and Environmental Research under contract DE-AC02-06CH11 357. The experiments at SACLA were performed at BL2 with the approval of the Japan Synchrotron Radiation Research Institute (JASRI) (Proposal No. 2018A8055 and 2019A8007). EAS was supported by NSF-MCB-RUI 1413360, NSF-MCB-EAGER grant 1839513 and NSF STC BioXFEL center award 6227. MN and LA training was supported in part by the National Institute of General Medical Sciences (NIGMS) of the National Institutes of Health (NIH) Maximizing Access to Research Careers (MARC) -T34 GM105549 grant. Use of BioCARS was supported by NIH NIGMS under grant number P41 GM118217. This research is partially supported by Platform Project for Supporting Drug Discovery and Life Science Research (Basis for Supporting Innovative Drug Discovery and Life Science Research (BINDS)) from Japan Agency for Medical Research and Development (AMED)). Structural models as well as structure factor amplitudes were deposited in the Protein Data Bank with accession codes 7JR5 and 7JRI for the wild-type 5ns and 33 ms structures of *Sa*BphP2 PCM, respectively.

## Author Contributions

Conceptualization, E.A.S. and M.S.; Methodology, S.W., E.A.S. and M.S.; Samples, M.C., J.S., M.N., L.A., D.F. and E.A.S.; Data Collection, M.C., S.P., J.S., M.N., I.P., L.A., T.N.M., E. C., W. Y. W., D. F., V. Š., M.M., L.C., S.I., E.N., R.T., T.T., L.F., K.T., S.O., S.W., E.A.S. and M.S.; Data Processing: I.P., L.C. and S.P.; Data Analysis, S.P., E.A.S. and M.S.; Writing – Original Draft, E.A.S and M.S.; Writing – Review & Editing, E.A.S. and M.S. with input from all authors.; Funding Acquisition, S.I., S.W., E.A.S. and M.S.; Resources and Supervision, S.I., S.W., E.A.S. and M.S..

## Corresponding Authors

Sebastian Westenhoff, Emina Stojkovic, Marius Schmidt

## Declaration of Interests

The authors declare no competing interests.

## Extended Data Figures and Tables

**Extended Data Figure 1.**
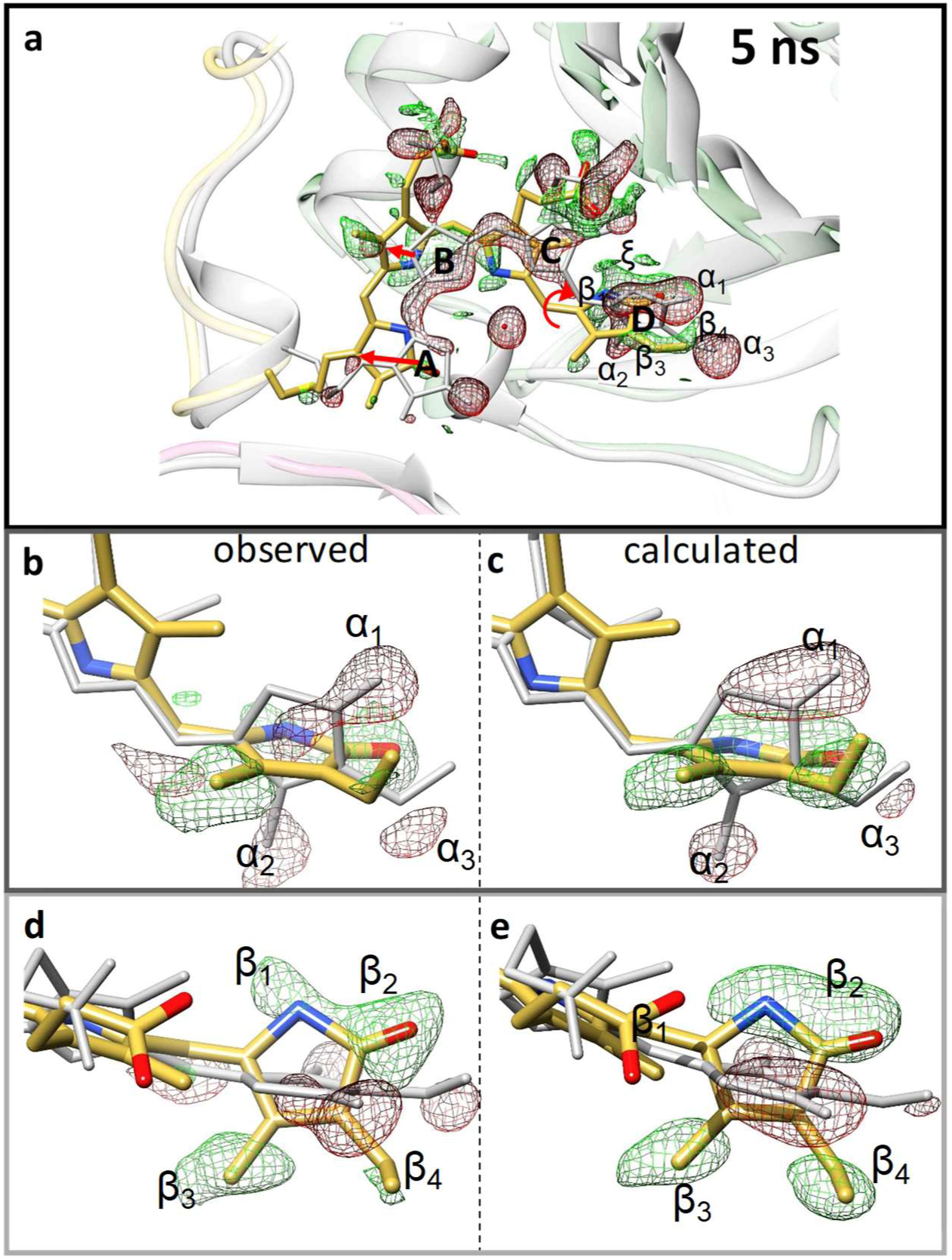
Chromophore isomerization 5 ns after excitation in subunit. **A**. DED is displayed on contour levels: red, −2.7 σ (- 3 σ of DED_66ms_), green, 2.7 σ (3 σ of DED_66ms_). (a) overall chromophore configuration. Gray: reference, dark, yellow: early intermediate at 5 ns. Red arrows show the direction of the chromophore sliding, and the direction of the rotation. (b) BV D-ring enlarged, side view as in (a) to emphasize the negative DED features, (c) corresponding calculated DED on the 4 σ contour level. (d) top view to emphasize positive DED features, and (e) corresponding calculated DED. The observed negative features α_1_ – α_3_ and the positive features β_1_ – β_4_ are reproduced, features ξ not shown in b – e for clarity.

**Extended Data Figure 2.**
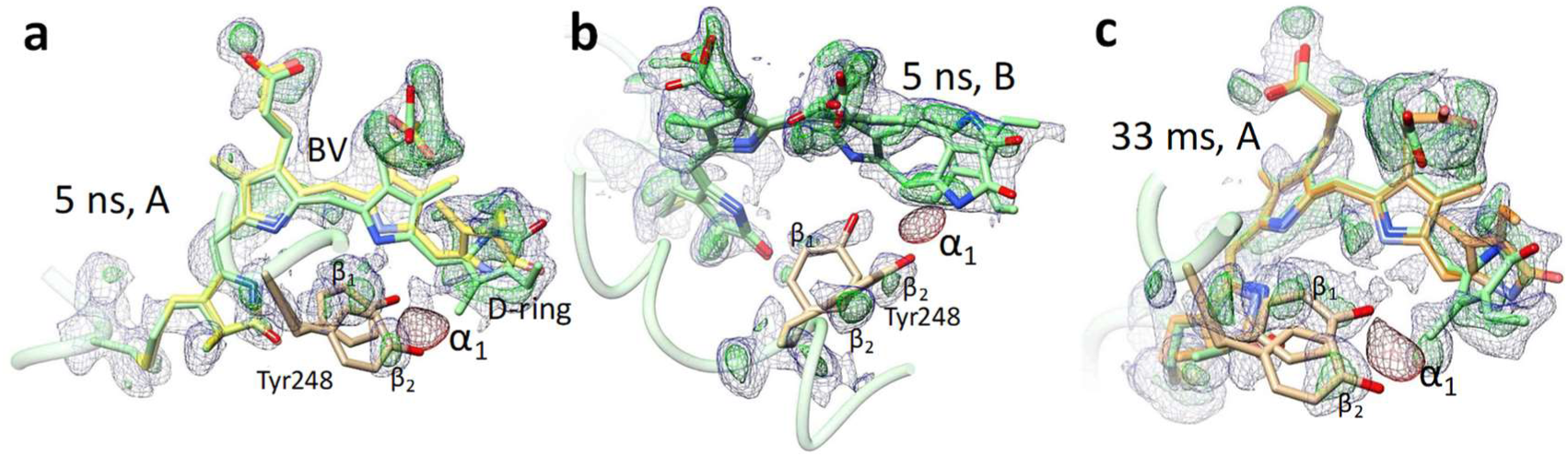
Close views of the BV chromophore and Tyr-248. Red and green maps: DED map (-/+ 2.7 σ contour level). Faint blue maps: extrapolated maps (N=19 and 22, for 5ns and 33 ms, respectively, contour level 1.5 σ). Only the negative feature α_1_ is displayed to avoid overcrowding as the reference structure is not shown. α_1_ is occupied by the Tyr248 hydroxyl of the reference. BV and Tyr248 are marked. (a) 5ns, subunit A. Chromophore in green: 90° ring-D rotation, clockwise; yellow: *E*-configuration. (b) 5ns, subunit B: double conformation of ring-D, clockwise and anticlockwise rotation. (c) 33 ms, subunit A. Green and gold chromophore conformations as in (a), respectively. Multiple positive features β indicate a double conformation of Tyr248 (β_1_ and β_2_, respectively) which is corroborated by the extrapolated maps.

**Extended Data Figure 3.**
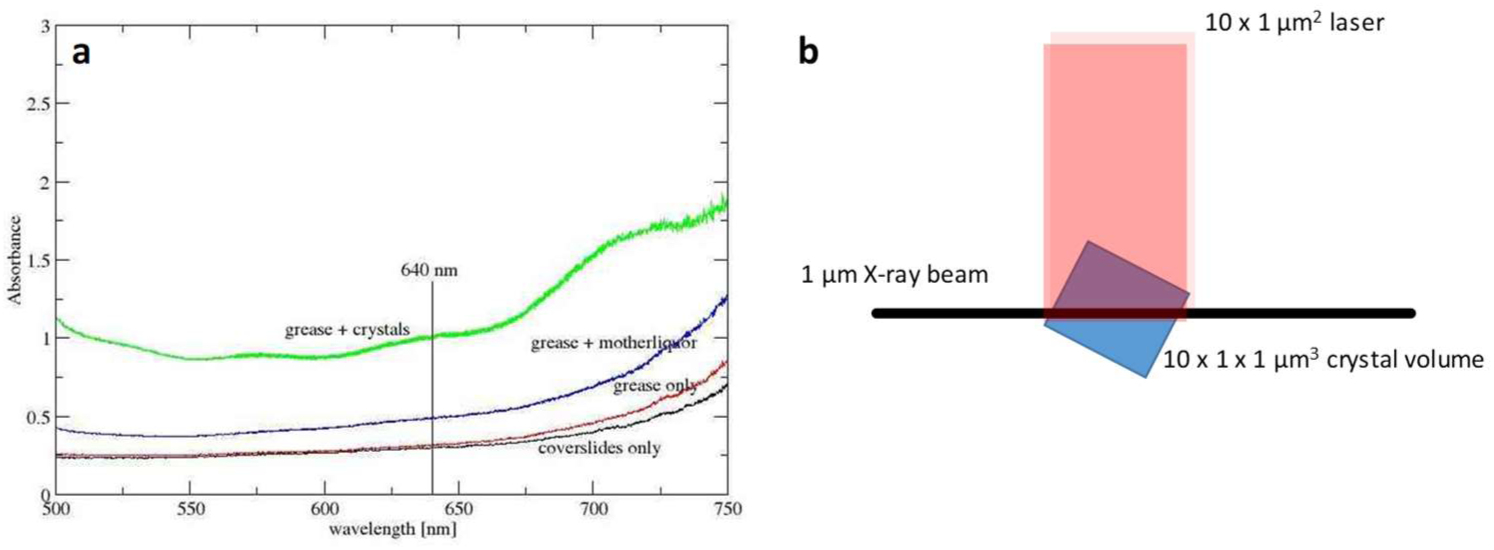
Absorption measurement on crystal-grease mixtures and perpendicular beam geometry. (a) At 640 nm (the excitation wavelength, marked) the grease is almost free of absorption (red line). With an optical path of 50 μm, coverslides, grease and crystals result in about an absorption of 1 at 640 nm (green line). (b) The effective (probed) illuminated crystal volume is given by the X-ray beam cross-section and the crystal size. The effective laser cross section is given by the crystal size and the X-ray beam diameter. The situation is depicted here with a 10 μm crystal and a 1 μm^2^ X-ray beam cross-section.

**Extended Data Figure 4.**
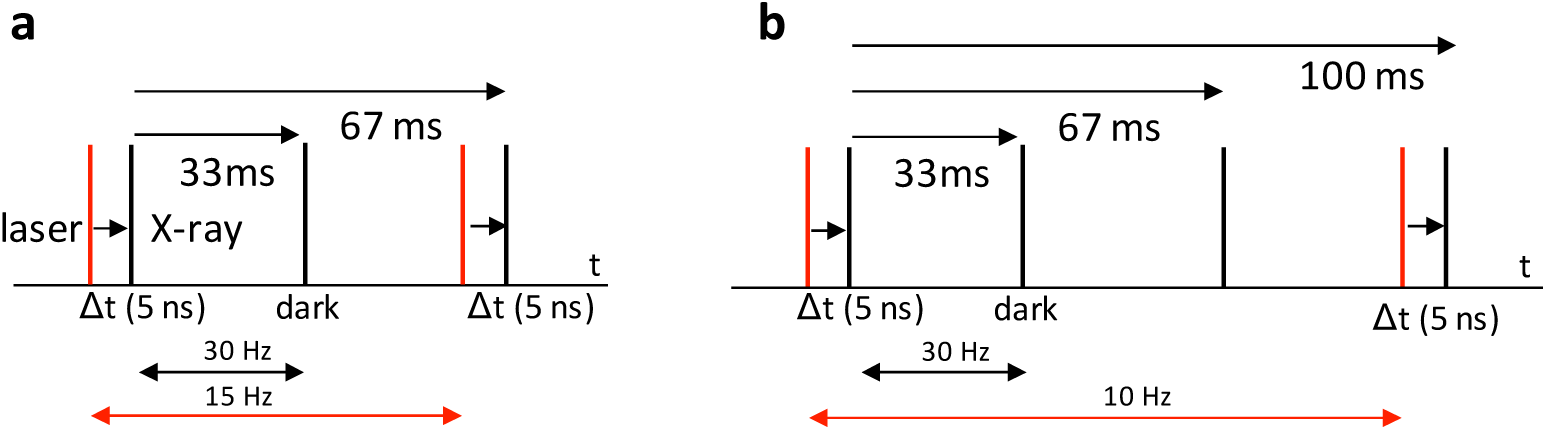
Data collection schemes. (a) Pump-probe, dark, pump-probe dark sequence. 30 Hz X-ray pulse repetition. The pump laser repetition rate is 15 Hz with 66 ms between laser excitations (b) Pump-probe, dark, dark, data collection sequence with a 10 Hz laser repetition rate with 100 ms between excitations.

**Extended Data Figure 5.**
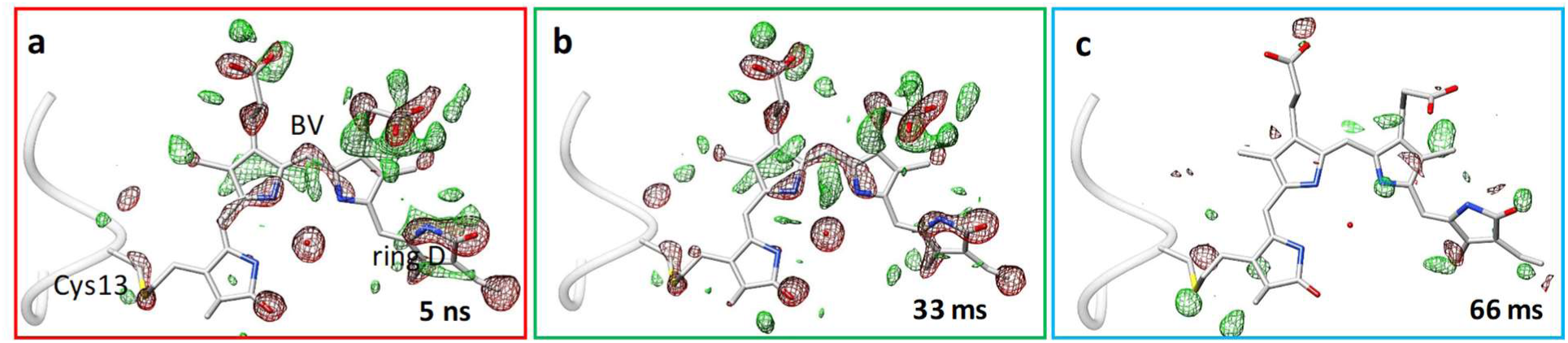
Difference maps near the chromophore. 5 ns (a), 33 ms (b) and 66 ms (c) after laser excitation (the reference structure is shown in gray). Green and red DED features are contoured at 2.7/-2.7 sigma for (a) and (b) and at 2.5/-2.5 sigma for (c), respectively. As there is signal in (a) and (b), at 66 ms the signal has vanished, and only spurious, random features persist.

**Extended Data Figure 6.**
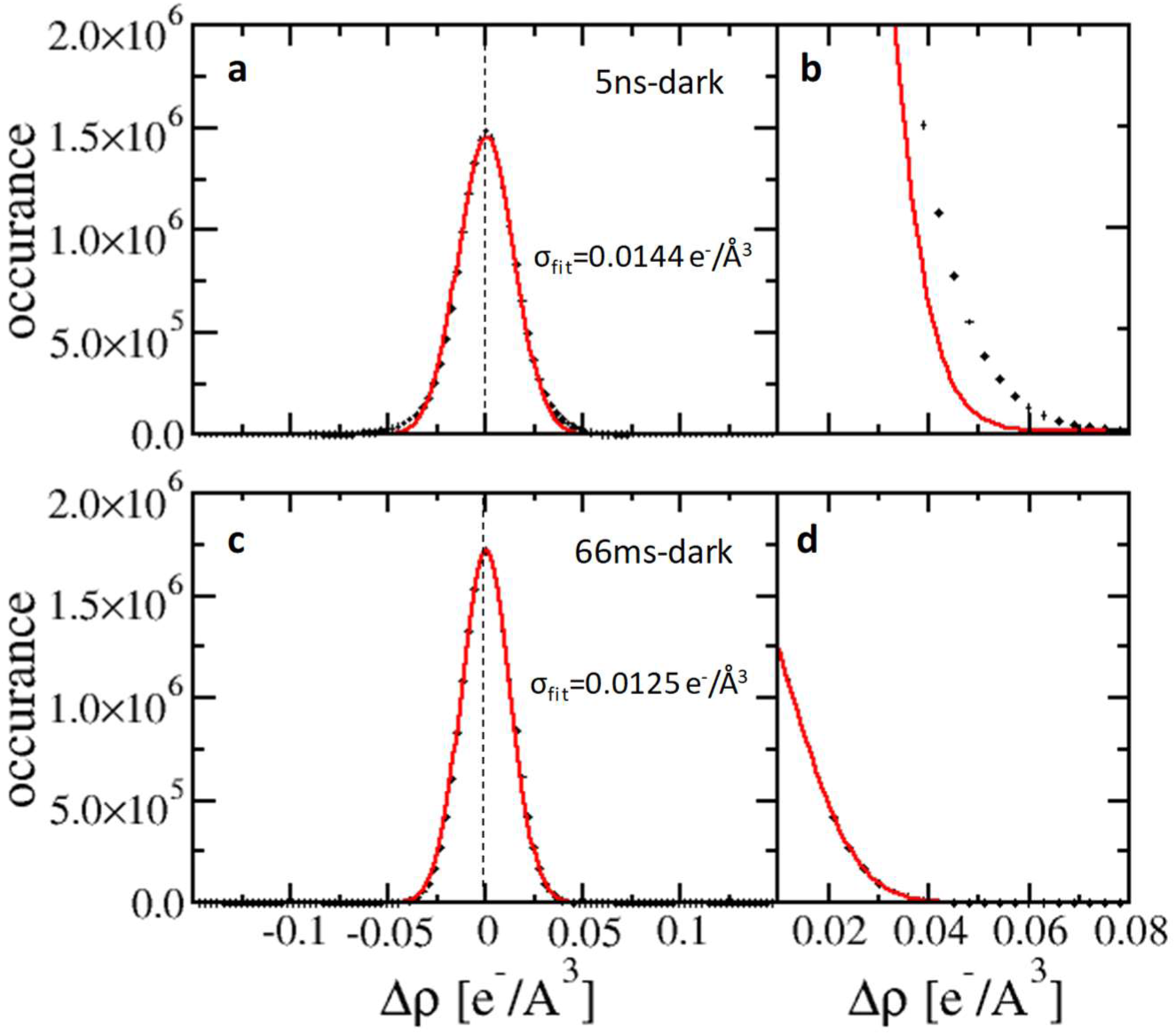
Histograms from DED maps. (a) and (b) Histograms derived from DED maps at 5ns. The Gaussian fit is shown in red. (b) enlargement of the fit of the Gaussian at the flanks of the histogram. (c) and (d) Histogram derived from the 66 ms – dark DED map. The fit of a Gaussian is perfect also in the flanks (d).

**Extended Data Figure 7.**
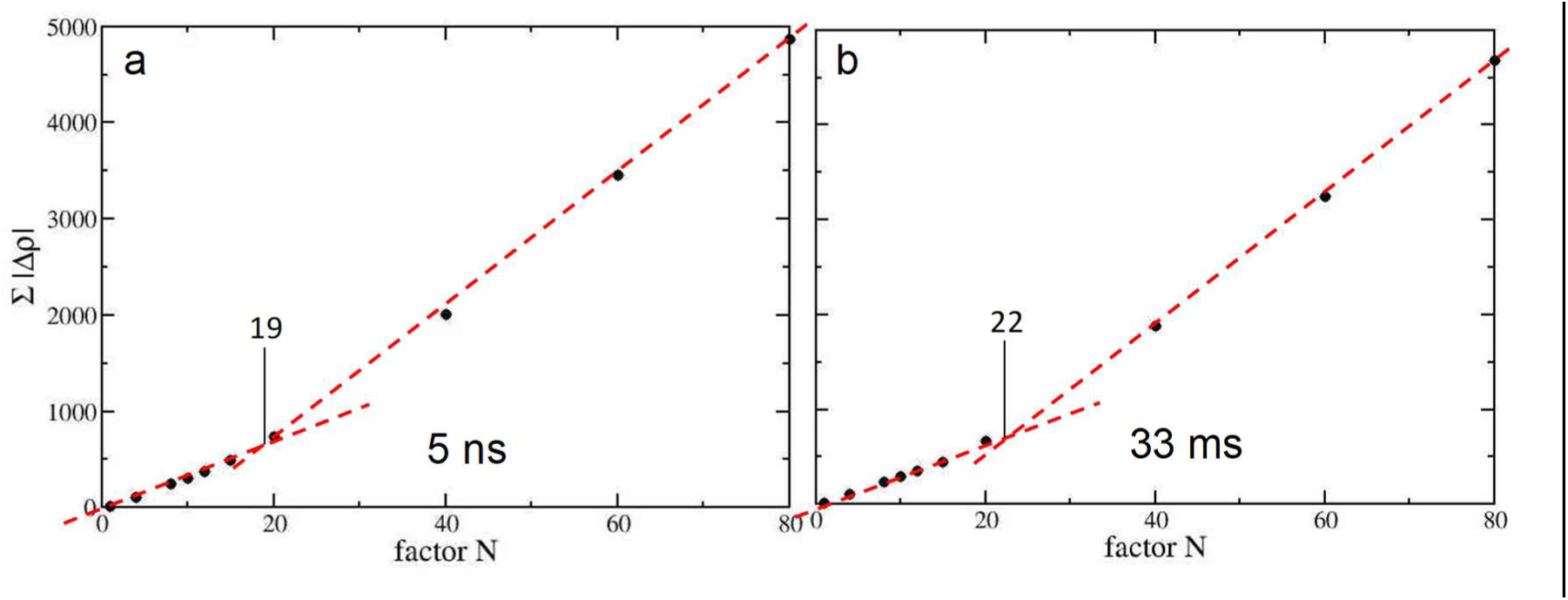
Factor N determination for extrapolated maps. The negative density ∑ |Δ*ρ*| found in extrapolated maps in a sphere of 6 Å about the ring D double bond is plotted against N. (a) The characteristic N (N_C_) for 5 ns is 19, (b) N_C_ is 22 for the 33 ms time delay.

**Extended Data Figure 8.**
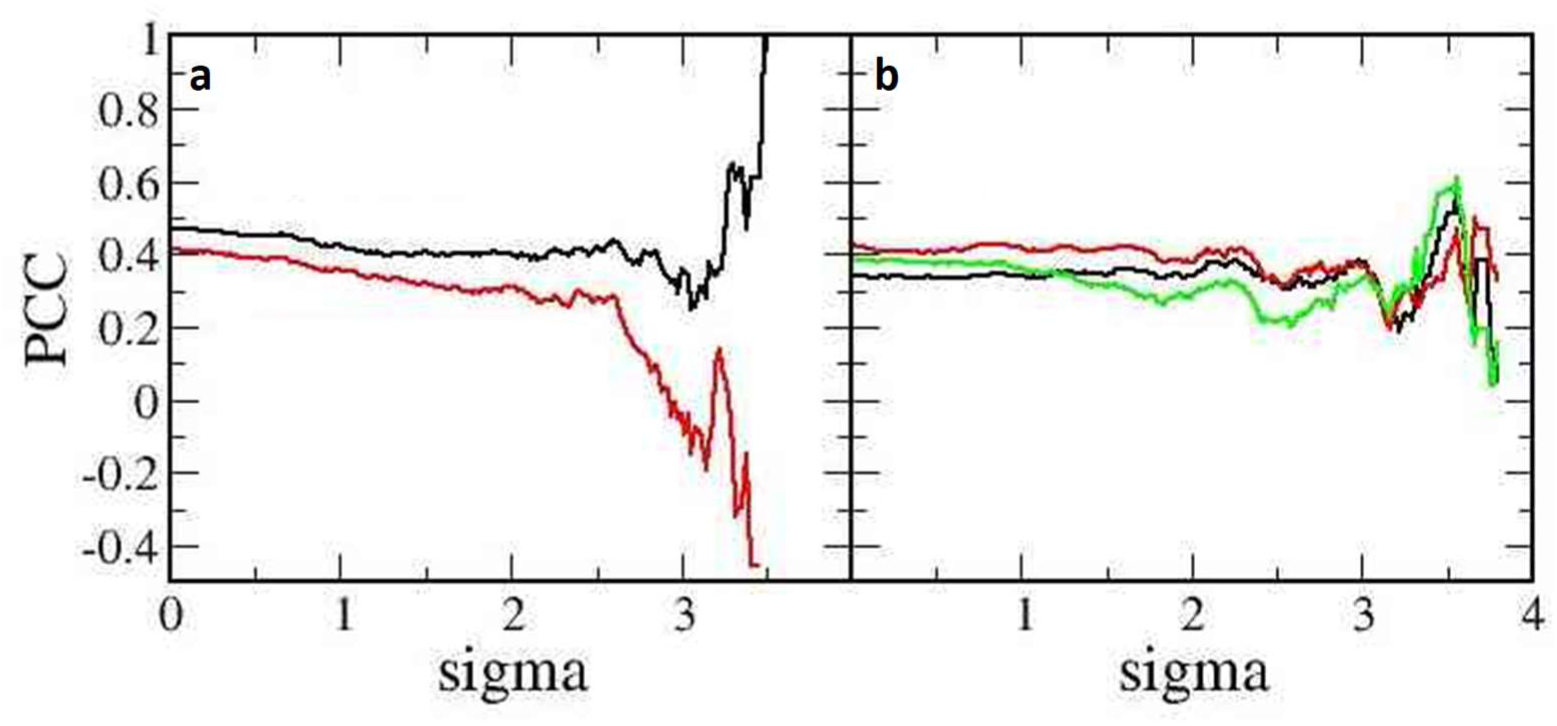
Pearson correlation coefficients as obtained by various models. (a) Subunit A, black line: ring-D clockwise rotation, red line: ring-D anticlockwise rotation. (b) Subunit B: black and red as in (a), green: double conformation.

**Extended Data Table 1.** Data collection statistics.

|  | Dark | 5 ns | 33 ms | 66 ms |
| --- | --- | --- | --- | --- |
| Beamline | SACLA BL2 |  |  |  |
| Resolution | 2.1 Å |  |  | 2.3 Å |
| Temperature | 285 K |  |  |  |
| Space group | P2<sub>1</sub> |  |  |  |
| Unit-cell parameters (°,) | a = 83.7 Å b = 83.4 Å c=86.9 Å $\alpha=90^\circ$ $\beta=107.6^\circ$ $\gamma=90^\circ$ | | | |
| Observations | 35,170,843 | 30,128,819 | 30,917,561 | 10,193,417 |
| No of unique reflections | 68,911 | 68,919 | 68,919 | 53,717 |
| Redundancy | 510 (111) | 437 (135) | 449 (154) | 190 (124) |
| Completeness (%) | 100.0 | 100.0 | 100 | 100 |
| CC1/2 at d<sub>min</sub> | 0.11 | 0.16 | 0.21 | 0.31 |
| R<sub>split</sub> (%) | 11.1 (221.9) | 9.4 (109.4) | 9.4 (101.6) | 18.1 (157.2) |
| Max/min $\Delta\rho$ | reference | 7.0/-8.6 | 6.7/-8.6 | 5.2/-4.9 |
| R<sub>cryst</sub>/R<sub>free</sub> (%) | 19.5/23.7 | 31.4/34.1 | 29.3/33.3 | -- |
| D-ring torsion | 32°/26° | A: -60°/-151°<br>B: 134°/-88° | A: -52°/-160°<br>B: 132°/-100° | -- |
| RMSD to reference [Å] | n/a | 1.4 | 1.5 | -- |

**Extended Data Table 2.** Difference electron density features and displacements. Values are listed for selected functional groups and atoms at the 5 ns time-delay. I applicable, displacements are averaged over double conformations.

| | Prominent DED Features, 5 ns | | Displacements [$\text{\AA}$] | |
| --- | --- | --- | --- | --- |
|  | Subunit A<br>Negative/positive<br>[sigma] | Subunit B<br>Negative/positive<br>[sigma] | 5ns |  |
|  |  |  | Subunit A | Subunit B |
| A-ring carbonyl | -5.5 / 3.5 | -7.3 / n.o. | 1.3 | 1.6 |
| B-ring methyl | -3.4/2.8 | -3.3/4.3 | 1.4 | 1.5 |
| B-ring nitrogen | -4.5/3.7 | -4.0/4.0 | 0.7 | 1.2 |
| B-ring propionate<br>carboxyl | -5.4/4.6 | -5.4/5.0 | 1.5 | 1.5 |
| C-ring propionate<br>carboxyl | -6.1/3.9 | -7.0/7.0 | 1.3 | 1.3 |
| D-ring carbonyl | -8.5/3.8 | -6.6/3.9 | 2.0 | 1.8 |
| D-ring methyl | -4.0/3.4 | -5.7/4.3 | 3.5 | 2.4 |
| D-ring vinyl<br>(double bond) | -3.3/3.2 | -3.7/2.5 | 3.1 | 2.2 |
| D-ring nitrogen | -5.3/3.2 | -3.1/3.5 | 3.4 | 2.3 |
| Tyr-201-OH | -6.0/4.2 | -5.8/4.8 | 1.4 | 1.3 |
| Tyr-248-OH | -5.5/4.1 | -5.5/3.0 | 1.3 | 2.0 |
| Ser-257-OG | -5.9/4.2 | -6.1/3.7 | 1.6 | 1.4 |
| Ser-259-OG | -4.6/4.0 | -5.0/3.5 | 1.2 | 2.5 |
| Ser-261-OG | -5.4/3.3 | -6.3/4.0 | 1.8 | 1.0 |
| Pyrrole water* | -6.1 | -5.4 | ex | ex |

\* photo-dissociated from binding pocket
n/o: not observed (positive and negative DED cancel)

**Extended Data Table 3.** Observed DED features and expected random DED features. Random features were estimated as a function of a multiple (x) of sigma (sigma level) in the unit cell of the *Sa*BphP2 difference map calculated on a 88 x 88 x 88 grid (N = 681472). The number of observed peaks was determined by the ccp4 program *peakmax* as a function of the DED sigma value. PM is the probability to observe at least one random feature equal or more than a given sigma level in the *entire* map, as opposed to the probability PI to observe this random feature in an individual voxel. Note, we write (1 – erf) rather than using the equivalent erfc, where erf is the error-function.

| Feature [sigma] | Observed | $\text{erf}(x/\sqrt{2})$ | PI<br>$1 - \text{erf}(x/\sqrt{2})$ | $\text{erf}(x/\sqrt{2})^N$ | PM<br>$1 - \text{erf}(x/\sqrt{2})^N$ | expected random features in the DED map<br>$N * (1.0 - \text{erf}(x/\sqrt{2}))$ |
| --- | --- | --- | --- | --- | --- | --- |
| 1 | > 8000 | 0.68 | 0.31 | 0.0 | 1.0 | 216238 |
| 2 | > 8000 | 0.95 | 0.05 | 0.0 | 1.0 | 31007 |
| 3 | 3167 | 0.997 | 0.002 | 0.0 | 1.0 | 1839 |
| 4 | 1018 | 1.0 | 0.0 | 0.0 | 1.0 | 43 |
| 5 | 264 | 1.0 | 0.0 | 0.67 | 0.33 | 0.4 |
| 6 | 44 | 1.0 | 0.0 | 1.0 | 0.0 | 0.0 |

**Extended Data Table 4.** Comparison of the reference (dark) and the 5 ns and the 33 ms *Sa*BphP2 structures. Bold entries: important interactions. Bold numbers: large changes relative to the reference structure.

|  | reference (Pr) |  | 5 ns |  | 33ms |  |
| --- | --- | --- | --- | --- | --- | --- |
|  | A | B | A | B | A | B |
| Asp192-O/A-ring N | 3.0 | 3.3 | 3.1/3.1 | 4.4/4.0 | **3.8/4.0** | **4.1/4.6** |
| Asp192-O/A-ring carbonyl | 4.5 | 4.4 | 4.1/4.1 | 5.5/5.8 | 4.5/5.1 | 5.4/5.6 |
| **Asp192-OD2/A-ring carbonyl** | 3.2 | 3.2 | **4.1/4.1** | **4.1/4.1** | **3.4/4.6** | **3.9/4.1** |
| Asp192-O/B-ring N | 3.0 | 3.0 | 2.9/3.0 | 3.1/3.3 | **4.2/4.3** | **3.3/3.3** |
| Asp192-O/C-ring N | 2.9 | 3.0 | **3.8/4.1** | **5.1/5.2** | **5.1/5.1** | **4.7/5.0** |
| **Asp192-OD2/Arg457-NH** | 2.6 | 2.9 | **5.4** | 3.0 | **4.4** | **4.8** |
| Asp192-OD2/Tyr161-OH | 5.6 | 5.7 | 5.5 | 4.9 | 5.6 | 4.2 |
| **Asp192-OD2/Tyr248-OH** | 2.7 | 2.7 | **3.5/3.5** | **5.8/5.3** | **5.3/5.5** | **3.9/4.4** |
| Tyr248-OH/D-ring methyl | 3.5 | 3.5 | 2.7/4.2 | 3.4/>5.0 | 2.6/3.9 | 2.9/5.3 |
| Tyr248-OH/Arg457-NH1 | 4.2 | 4.4 | 3.5/>**5.0** | 3.5/4.3 | 5.3/5.5 | 4.8/5.5 |
| **Ser242-N/B-ring propionyl O2** | 2.9 | 2.8 | **5.1/5.1** | **4.3/>5.0** | **3.3/3.3** | **5.2/5.7** |
| **Ser257-OG/C-ring propionyl O2** | 3.2 | 2.5 | 2.3/2.6 | 2.6/**3.9** | 2.6/**3.3** | **3.5/5.0** |
| **Ser259-OG/C-ring propionyl O2** | 2.7 | 2.7 | 3.1/3.3 | 2.6/2.5 | 2.1/**3.5** | 2.3/2.4 |
| Ser259-OG/Arg207-NH2 | 2.9 | 2.8 | **5.1** | **4.7** | **4.4** | **5.4** |
| **His275-NE2/C-ring propionyl O2** | 4.7 | 6.2 | 6.0 | 7.1/7.2 | 6.7/6.9 | 6.2/7.5 |
| **His275-NE2/D-ring carbonyl** | 3.0 | 2.8 | **6.0** | 2.8/**4.3** | **4.3/4.8** | **3.3/4.4** |
| His275-ND1/Arg157-NH2 | 2.9 | 2.9 | **3.6** | **3.8** | 2.7/**4.8** | **4.5** |
| His245-ND1/B-ring N | 3.5 | 3.6 | 3.6/3.8 | 3.7/3.8 | 3.7/3.7 | 5.1/5.3 |
| His245-ND1/C-ring N | 3.4 | 3.6 | 3.7/3.8 | 4.2/4.7 | 3.7/3.8 | 4.6/5.4 |
| **His245-NE2/C-ring propionyl O1** | 2.6 | 2.8 | 2.7 | 2.9/**3.8** | 3.0/**3.5** | **3.4/5.4** |
| Arg457-NH1/A-ring carbonyl | 4.5 | 4.5 | 6.0 | 5.4/5.5 | 7.4/7.4 | 7.0/7.2 |
| Tyr201-OH/Arg207-NH1 | 4.0 | 3.9 | 3.7 | 4.0 | 4.9 | **3.0** |
| Tyr201-OH/B-ring propionyl O1 | 2.7 | 2.8 | 2.9 | 2.7/3.1 | 2.8/2.9 | 2.3/3.1 |
| Tyr201-OH/C-ring propionyl O1 | 5.9 | 5.9 | 4.4/5.9 | 5.0/5.0 | 5.9/6.1 | 5.7/5.7 |
| Ser261-OG/ Arg207-NE | 3.1 | 2.7 | **3.6** | **4.0** | **3.6** | **4.0** |
| Arg207-NH2/C-ring propionyl O1 | 5.4 | 5.3 | 4.9/6.1 | 4.1/5.7 | 4.1/5.1 | 3.9/4.2 |
| Arg239-NH2/B-ring propionyl O1 | 2.7 | 2.9 | 2.4 | 2.5/2.6 | **3.8/3.9** | 2.8/2.9 |
| A-Ring torsion | 7 | 9 | 12/20 | 40/40 | 7/-1 | 15/39 |
| D-Ring torsion | 32 | 27 | -61/-151 | 134/-88 | -62/-160 | 132/-100 |
| B/C-Ring relative tilt* | 11 | 14 | 25/25 | 43/37 | 32/46 | 43/46 |
| Δ D-ring rotation** | - | - | -93/<br>**-183** | 102/<br>-120 | -94/<br>**-192** | 105/<br>-127 |

\* separately for the two alternate conformations
\*\*difference between D-ring torsion in the dark and the D-ring torsion at measured time delays. **Negative** values: clockwise rotation relative to dark, **positive** values: counterclockwise rotation relative to dark, bold red: *E*-configuration.

**Extended Data Table 5.** Important distances in other BphP structures. (identified by pdb entry). Sequence numbers are in the *Sa*BphP2 PCM convention (directly comparable with Extended Data Tab. 4), they denote equivalent amino acid residues found in the various structures. L3 is an intermediate determined by temperature scan crystallography.

|  | 1ps<br>6TSU |  | *DrBphP* (Pfr)<br>5C5K |  | Bathy *PaBphP*<br>(Pfr)<br>3C2W |  | L3<br>(bathy)<br>3NOU |
| --- | --- | --- | --- | --- | --- | --- | --- |
|  | A | B | A | B | A | B | A |
| Asp192-O/A-ring N | 2.7 | 3.2 | 2.8 | 2.8 | 3.2 | 3.1 | 3.0 |
| Asp192-OD2/A-ring carbonyl | 4.3 | 4.8 | 4.1 | 4.2 | 3.5 | 2.8 | 3.3 |
| Asp 192-O/A-ring carbonyl | 3.8 | 3.8 | 3.9 | 3.6 | 4.1 | 3.4 | 3.9 |
| Asp192-O/B-ring N | 3.0 | 3.4 | 3.1 | 3.3 | 3.1 | 3.6 | 2.7 |
| Asp192-O/C-ring N | 3.4 | 3.5 | 3.5 | 3.4 | 3.0 | 3.3 | 2.7 |
| Asp192-OD2/Arg457-NH | n/a | n/a | 16.1 | 15.8 | nd | nd | nd |
| **Asp192-OD2/Tyr248-OH** | 5.9 | 5.2 | 3.8 | 3.8 | 4.0 | 4.5 | 3.9 |
| Tyr248-OH/D-ring carbonyl | 4.1 | 4.0 | 3.5 | 3.1 | 3.2 | 3.9 | N/A |
| **Ser242-N/B-ring propionyl O2** | 3.2 | 3.3 | 4.9 | 4.7 | 5.1 | 4.8 | 5.5 |
| Ser257-OG/C-ring propionyl O2 | 3.7 | 4.0 | 3.1 | 2.2 | 2.5 | 3.5 | 4.7 |
| **Ser259-OG/C-ring propionyl O2** | 4.5 | 3.6 | 6.8 | 3.9 | 6.7 | 6.2 | 2.5 |
| Ser259-OG/Arg207-NH2 | 3.2 | 3.7 | 7.5 | 7.7 | nd | nd | nd |
| **His275-NE2/C-ring propionyl O2** | 5.4 | 5.7 | 3.8 | 5.0 | 2.8 | 3.4 | 4.1 |
| His275-NE2/D-ring carbonyl | 3.9 | 4.3 | 7.7 | 8.6 | nd | nd | 6.6 |
| **His275-ND1/Arg157-NH2** | 4.9 | 4.3 | 6.9 | 6.5 | 4.7 | 4.2 | 4.8 |
| His245-ND1/B-ring N | 3.6 | 3.5 | 3.6 | 4.5 | 3.7 | 3.7 | 4.2 |
| His245-ND1/C-ring N | 4.0 | 4.2 | 3.6 | 3.7 | 3.8 | 3.9 | 4.3 |
| **His245-NE2/C-ring propionyl O1** | 2.8 | 2.7 | 2.8 | 3.7 | 4.2 | 4.2 | 5.6 |
| Tyr201-OH/Arg207-NH | 4.7 | 4.4 | 7.5 | 7.2 | nd | nd | nd |
| Tyr201-OH/B-ring propionyl O1 | 2.7 | 3.8 | 3.8 | 3.1 | 3.8 | 2.8 | 2.8 |
| **Tyr201-OH/C-ring propionyl O1** | 7.3 | 6.0 | 5.7 | 3.3 | 5.4 | 5.1 | 2.9 |
| Ser261-OG/ Arg207-NH1 | 2.7 | 2.8 | nd | nd | nd | nd | nd |
| Arg239-NH2/B-ring propionyl O1 | 3.4 | 3.0 | 3.5 | 2.7 | 4.0 | 4.1 | 4.6 |
| Ring-A torsion | 19 | 40 | -7 | 10 | 11 | 30 | 24 |
| Ring-D torsion | 58 | 89 | -142 | -161 | -150 | -143 | -25*** |
| B/C-Ring relative tilt | 8 | 9 | 0 | 0 | 5 | 6 | 35 |
| Δ ring D rotation | 33* | 69* | -179** | -198** | na | na | 125 |

\*difference from comparing pdb entries 6TSU/6T31, **counterclockwise** rotation (positive)
\*\* difference from comparing pdb entries 5C5K/4Q0J, **clockwise** rotation (negative)
\*\*\* bathy phytochrome: -150° in the dark, rotation relative to dark is **counterclockwise** (positive)
nd: not done, n/a not applicable because the residue is missing.

## Notes

### Competing Interest Statement

The authors have declared no competing interest.

